# Tropomyosin and Vinculin antagonistically mediate GPI-anchored protein nanoclustering

**DOI:** 10.64898/2026.04.30.721936

**Authors:** Sarayu Beri, Teresa Massam-Wu, Satyajit Mayor

## Abstract

The plasma membrane is laterally organised into dynamic nanoscale and mesoscale domains. Glycosylphosphatidylinositol-anchored proteins (GPI-APs) form actomyosin dependent nanoclusters at the outer leaflet of the plasma membrane via transbilayer coupling to inner leaflet lipids. These nanoclusters are nucleated downstream of integrin activation in a mechanochemical fashion and require vinculin and myosin 1 activity. Here, we identify an antagonistic relationship between vinculin and a negative regulator of myosin 1, tropomyosin, in regulating GPI-AP nanoclustering. We show vinculin mediated restoration of clustering is myosin-1 sensitive. The actin and lipid-binding vinculin tail is sufficient to restore clustering, indicating that vinculin, in particular its actin and lipid-proximal tail promote nanoclustering. Furthermore, re-expression of the non-muscle tropomyosin isoform Tpm2.1 in Tpm2-deficient MDA-MB-231 cells suppress GPI-AP nanoclustering while Tpm2.1 mutants defective in stable actin-filament association do not. Structural superposition of actin-bound vinculin tail and tropomyosin assemblies suggests that vinculin and tropomyosin engage overlapping or closely adjacent surfaces on actin filaments. Consistent with this, depletion of tropomyosin in vinculin-null cells restores GPI-AP nanoclustering. Together these results suggest that competitive interactions between tropomyosin and vinculin regulate actin availability for myosin 1 and hence contribute to the regulation of GPI-AP nanoclusters.

## Introduction

Molecular constituents of the fluid bilayer of the living plasma membrane are laterally heterogenously distributed in a regulated manner ^1–3^. Membrane proteins and lipids form nano and mesoscale domains that influence membrane-associated cellular processes such as signalling and trafficking^1,4–8^.For example, the construction of focal adhesion complexes determine how cells sense the external environment of the cell, and the construction of endocytic pits determine how membrane proteins and nutrients are internalized into the cell^9,10^. Understanding how these distinct membrane domains are generated and regulated is therefore important for understanding how membrane structure supports cellular function. Along with passive mechanisms of clustering mediated by specific binding interactions dictated by thermodynamics^5^, active mechanisms, in particular interaction of the membrane with active stresses derived from cortical actomyosin have been deployed by the cell to regulate the structure and architecture of membrane domains^11–14^.

Recent work from our laboratory has shown that different classes of membrane proteins depend on distinct actomyosin complexes for clustering^15^. In particular, the lipid binding Class I non-muscle myosin motor family are required for clustering of Glycosyl-phosphatidylinositol-anchored proteins (GPI-APs), while the cortically localized Class II non-muscle myosin motors are involved in clustering of actin binding transmembrane proteins. Thus, the physical basis of clustering depends on the nature of the membrane-associated molecules as well as its coupling to a specific actin filament population and is therefore mechanistically and likely functionally specialized^15–17^.

In the context of the GPI-APs at the outer leaflet, it is likely that phosphatidylserine (PS) interacts with membrane-proximal actin filaments via phosphatidylinositol 4,5 bisphosphate (PIP_2_) and PS-binding motors such as myosin 1^18^. GPI-AP-nanoclustering necessarily involves transbilayer interactions between inner-leaflet immobilized PS and long, saturated acyl chains of the outer leaflet GPI-anchors^19^. Transbilayer interactions via interdigitation of the acyl-chains require cholesterol as a chain-straightening agent. This compositional configuration naturally leads to the generation of liquid-ordered (*lo)* nanodomains, and along with their lateral interactions, generate mesoscale active-emulsions with *lo*-like properties ^19–21^. Thus, actomyosin activity, coupled via PIP_2_ and PS, and transbilayer links with outer-leaflet GPI-APs configures the local composition and biophysical properties of the membrane in a dynamic manner.

The formation of mesoscale *lo* domains could serve as important signaling platforms. For instance, ordered domains act as hubs for lipid-modified small GTPases supporting the sustained activation of Rac1 GTPase responsible for lamellipodial extension during cell-spreading and migration^2,22–25^. Consistent with this, it was recently shown that GPI-AP-nanoclusters are generated downstream of integrin receptor signaling. Importantly, cells that were unable to generate GPI-AP-nanoclusters due to mutations in enzymes that prevent the attachment of long-acyl chains onto the GPI-APs during its biosynthesis ^26–28^, failed to create liquid-ordered domains and showed aberrant integrin-related-function such as spreading and migration defects^29^.

Since the nanocluster-rich domains are actively generated they are subject to dynamic regulation. In fact, integrin-engagement with the ligands in the extracellular matrix and under load, trigger the localized generation of nanocluster-rich domains, integrating chemical ligation, as well as mechanical inputs^29^. Fibronectin, an extracellular matrix protein, acts as a chemical ligand for the a_5_β_1_ containing integrin-receptor leading to the activation of RhoA, formins and myosin isoforms. The mechanical input is mediated by the mechanosensitive protein, talin and its mechanotransducer, vinculin. These proteins are recruited to both nascent and mature focal adhesions which are generated by the development of force at the integrin-ECM interface ^9^.

Vinculin is an important mechanotransducer that is involved in the formation of functional focal adhesions^9^. Through force-dependent activation downstream of talin, vinculin stabilises adhesions and transmits information about the mechanical environment to the cell and is necessary for directed cell locomotion in 2D and 3D environments^30 31,32^. Studies from our laboratory^29^ showed that vinculin is also required for GPI-AP nanoclustering: vinculin-null (Vin-/-) cells exhibit a loss of clustering, whereas re-expression of vinculin restores it. Clustering is mechanosensitive because when cells are plated on fluid lipid bilayers containing lipids conjugated with the integrin ligand RGD, they do not exhibit GPI-AP clustering downstream of integrin engagement. However, when integrins experience traction by encountering immobile RGD ligands in the fluid matrix, vinculin is recruited to the sites of integrin engagement resulting in localized nanoclustering.^29^ The mechanosensitive vinculin-talin interaction is necessary for vinculin activation, but once activated, vinculin can support clustering even in the absence of talin binding. However, the lipid and actin-binding activities of vinculin are strictly necessary for the nanoclustering process, and cannot be rescued even by using a constitutively activated lipid or actin non-binder. Thus, the plasma membrane organisation of GPI-APs is sensitive to mechanical cues and depends on vinculin’s interactions with lipids and actin. These observations together suggest that the membrane is dynamically configured and capable of building active emulsions in a spatially regulated manner. However, the molecular mechanism by which vinculin gates this process remains unclear.

Given the central role actomyosin activity plays in this process, one potential level of regulation is access of myosin 1 to actin filaments. Here, a negative regulator of myosin1, tropomyosins, are likely to play a central role. Tropomyosins are a family of actin-binding proteins that polymerise head-to-tail along actin filaments and regulate the binding of other actin-associated proteins^33–36^. Although tropomyosins are classically associated with muscle function, non-muscle isoforms are widely expressed and are increasingly recognised as important determinants of actin filament identity and accessibility. By coating actin filaments, tropomyosins can stabilize actin structures, alter filament accessibility, and govern the recruitment of motors and actin-binding proteins^37–43^. *In vitro* reconstitution studies have shown that non-muscle tropomyosin isoforms, including Tpm2.1, can inhibit myosin 1-actin interactions, suggesting that tropomyosin may regulate motor access to actin filaments available for membrane remodelling^37,39,44–47^.

Tpm2.1 is of particular interest in this context because it has been implicated in regulating mechanosensing in non-muscle cells. Altered Tpm2 expression has been associated with pathological states including cancer^48–50^. MDA-MB-231 cells which lack TPM2 expression^51^, provide a useful system in which to test the consequences of reintroducing Tpm2.1 on plasma membrane organisation. More broadly, since tropomyosins limit myosin 1 access to actin, then tropomyosin isoforms could function as a negative regulator of the actin-dependent membrane organisation required for GPI-AP nanoclustering.

Here we investigated how vinculin, and Tpm2.1 contribute to nanoclustering of GPI-APs. We first asked whether GPI-AP nanoclustering is sensitive to myosin 1 activity down-stream of vinculin activation. We then examined which vinculin functions are sufficient to support clustering, focusing on the actin- and lipid-binding properties of the vinculin tail domain. Finally, we tested whether Tpm2.1 negatively regulates nanoclustering and whether vinculin may promote clustering, at least in part, by reducing tropomyosin occupancy on actin and thereby increasing actin availability for myosin 1-dependent membrane organisation.

## Results

### Integrin-triggered GPI-AP nanoclustering requires the vinculin-tail domain

To assess the role of vinculin in nanoclustering of GPI-APs we utilized fluorescence emission anisotropy of fluorescently tagged GPI-APs to measure proximity between GPI-APs. Emission anisotropy reports on the extent of nanoclustering of GPI-APs; increased clustering is expected to enhance FRET between like fluorophore (homo-FRET) and therefore lower anisotropy^16,52^ (Fig. 1A,B). Using this technique, we have previously shown that vinculin is necessary for GPI-AP clustering down stream of integrin-mediated cell adhesion. When allowed to adhere to fibronectin-coated glass coverslips (hence activating integrin), vinculin knockout mouse embryonic fibroblasts (Vin-/-MEFs, hereafter Vin -/-), (Fig. 1C) exhibit a high GPI-AP anisotropy indicative of a lack of GPI-AP clustering (Fig. 1D), and they also spread poorly on fibronectin. (Fig. 1E). As shown previously^29^, reintroduction of vinculin restores the GPI-AP clustering as well as the ability of the cells to spread on fibronectin. This is disrupted when a cholesterol depletion agent, methyl-βcyclodextrin (MβCD) is added to the cells. Furthermore, treatment with a myosin 1-specific inhibitor pentachloropseudilin (PClP) ^53,54^ also leads to loss of GPI-AP nanoclustering^15^ in the Vin -/- cells when Vinculin expression was restored (Fig. 1 F-I). These results confirm a role for vinculin in promoting myosin 1 and cholesterol-dependent clustering of GPI-APs, downstream of integrin-activation and consequently cell spreading.

**Fig. 1:**
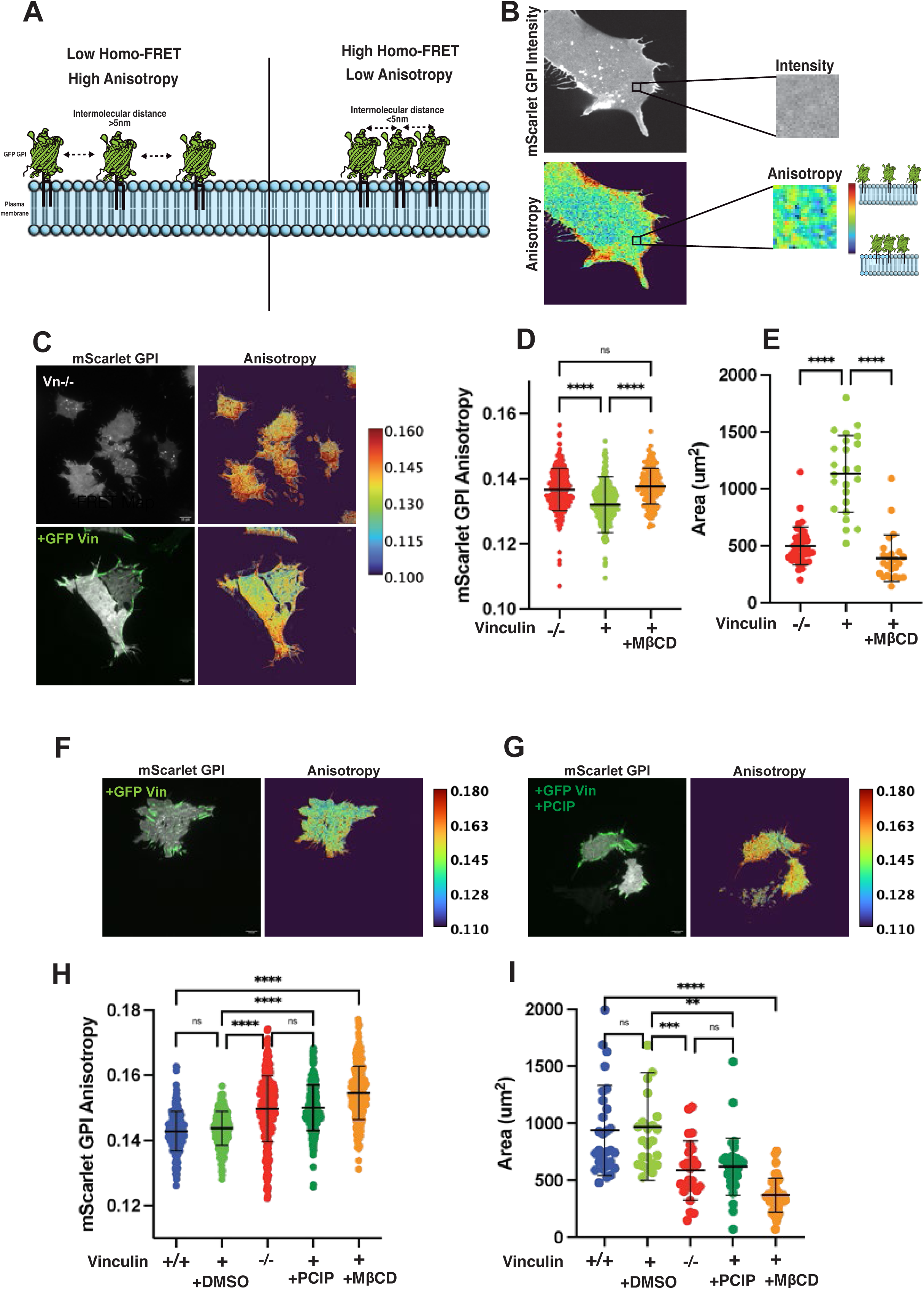
Vinculin regulates GPI-AP nanoclustering in a myosin1 dependent manner. A) Schematic depicts FRET between like fluorophores relating fluorescence emission anisotropy and homo-FRET. Fluorophores separated by distances greater than the Forster’s radius (eg.∼ 5nm for GFP) are unable to participate in energy transfer, resulting in high fluorescence emission anisotropy values while those within Forster’s radius participate in the FRET process, which consequently lowers emission anisotropy. B) Fluorescence intensity and emission anisotropy maps of a cell transfected with mScarlet-GPI. Insets show uniform fluorescence from a membrane patch, while the corresponding homo-FRET map highlights differences in anisotropy and the heterogeneity in the extent of homo-FRET between molecules in the patch. C) Fluorescence intensity (left panel) and anisotropy maps of mScarlet GPI expressed in cells lacking (Vin-/-) without or with co-expression of GFP-Vin (green). D, E) Scatter plots show mean ±s.d of emission anisotropy (D) determined from ROIs (2.2 x 2.2um^2^) such as those as defined in B, and Cell area (E) from cells treated as indicated. Data shown from representative experiment, N>3, Each condition from >22 cells F, G) Intensity images and anisotropy maps of mScarlet GPI expressed in with type MEFs (Vin +/+) or cells lacking Vin-/- co-expressing GFP Vin (green), without (F) or with myosin 1 inhibitor PClP treatment. H, I) Scatter plots of emission anisotropy (H) and Cell area (I) determined from cells as above, and treated as indicated. Data shown from representative experiment, N=3. Each condition from >21 cells. All intensity and anisotropy images were collected using TIRF emission anisotropy microscopy. Bars in the scatter plots represent mean ±s.d. *p* values determined from Kruskal-Wallis test with Dunn’s multiple comparisons test, where ns indicates no significant difference and **** indicates p <0.0001, *** <0.001 and **<0.0018. Scale bar, 10um

Previous studies have indicated that activation of talin is necessary for GPI-AP clustering, and mutants of vinculin that fail to bind talin are unable to support GPI-AP clustering ^29^. This is likely because vinculin is in an autoinhibited state wherein the vinculin head sequesters the actin and lipid-binding tail region; talin-mediated activation relieves this inhibition (Fig. 2A). A point mutation in the head region of vinculin which relieves the head-tail interaction leading to a constitutively active vinculin restores GPI-AP clustering even in the absence of talin binding. In contrast, actin and lipid-binding-deficient vinculin mutants fail to rescue clustering, and the head domain alone is insufficient^29^. Cell spread area measurements with various vinculin mutants show a corresponding dependence on lipid, actin and talin-binding capabilities of vinculin, consistent with the role of vinculin-mediated GPI-AP clustering on this process (Supplementary Fig. S1A). These observations suggest that the tail domain of vinculin may be the critical domain required for this process.

**Fig. 2:**
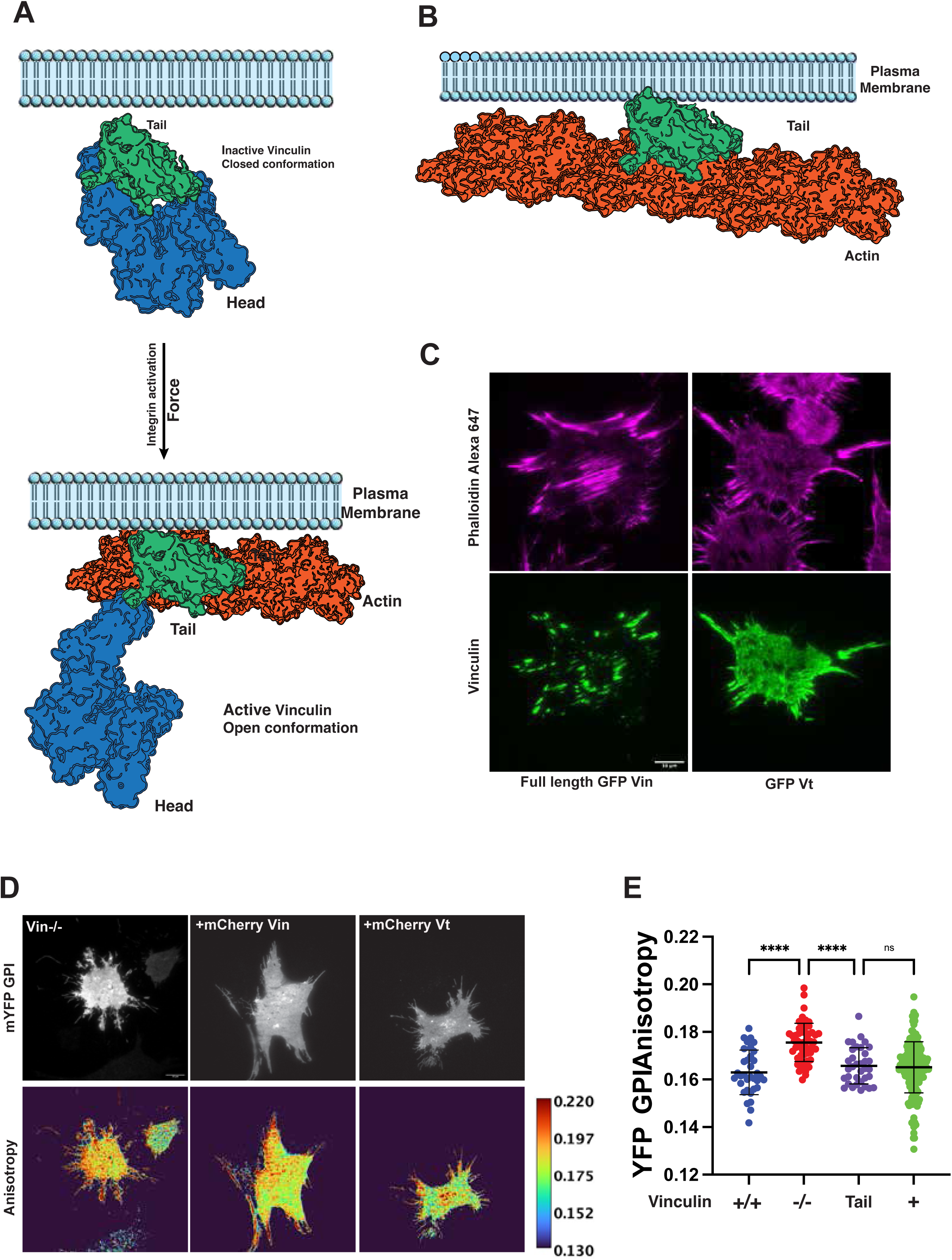
The vinculin tail domain is required for GPI-AP nanoclustering. A) Schematic depicts the autoinhibited (closed) and activated (open) conformation of vinculin. Force-mediated integrin activation resulting in relieving autoinhibition and exposing the vinculin tail domain for actin and lipid binding. B) Schematic depicts membrane and actin-association of the isolated vinculin tail domain representing a constitutively activated vinculin. C)TIRF images of full length GFP-tagged (green) vinculin and vinculin-tail expressing Vin-/- cells co-stained with Alexa 647-Phalloidin (magenta) marking actin filaments. D) Intensity images and anisotropy maps of mYFP-GPI expressed in cells lacking vinculin (Vin-/-) or co-expressed with mCherry-Vin or mCherry-Vin-t. E) Scatter plots show mean ±s.d of emission anisotropy determined from ROIs (1.4 x 1.4um^2^) from cells as indicated. Data shown from a representative experiment, N=2. Each condition from >16 cells. *p* values determined from Kruskal-Wallis test with Dunn’s multiple comparisons test, where ns indicates no significant difference and **** indicates *p* <0.0001. Scale bar, 10um

To test this directly, we expressed the vinculin tail domain (Vt), which contains residues required for both actin and lipid binding, in Vin -/- cells. In contrast to full-length vinculin, the isolated tail domain lacks the autoinhibitory head-tail interaction that normally masks the actin-binding site and therefore could potentially interact more constitutively with available actin filaments (Fig. 2B). This is corroborated by the extensive distribution of GFP-Vt on intracellular actin structures in contrast to the more restricted distribution of full length GFP-Vin to actin structures at focal adhesions (Fig. 2C). Transfection of mCherry-Vt in Vin-/- cells co-expressing mYFP-GPI resulted in lowering of mYFP emission anisotropy similar to that obtained by co-expression of mCherry-Vin^29^ (Fig. 2D, E), showing that Vt restored GPI-AP clustering. Expression of Vt also allowed cells to spread on fibronectin-coated glass, albeit not as effectively as wild type Vinculin (Supplementary Fig. S1). This may be due to the unregulated vinculin tail’s ability to act as a barbed end actin capping protein, preventing the polymerization of longer actin filaments required for cell spreading^55–57^. These data indicate that the vinculin tail domain alone is sufficient to support nanoclustering and suggest that vinculin promotes functional plasma membrane organization of GPI-APs via its ability to interact with actin at the plasma membrane.

### Tpm2.1 negatively regulates GPI-AP nanoclustering

Active nucleation of GPI-AP nanoclusters at the outer leaflet involves the transient immobilizing of PS at the inner leaflet by a linker that connects PS with actin. Considering that the class I non-muscle myosin 1 motors bind PIP_2_ and PS at the inner leaflet, it is likely that myosin 1 serves as a linker between actin and the inner leaflet-thus building a contractile platform necessary for this activity^58^. It is therefore surprising that nanoclustering is dependent on the presence and subsequent activation of vinculin. This is possible if vinculin counteracts negative regulators of myosin 1- actin interactions, such as tropomyosin. Support for this hypothesis stems from the observation that non-muscle isoforms of tropomyosin are abundantly present in all cells and are known to negatively regulate myosin 1 binding to actin^34^. *In vitro* reconstitution studies suggest that non-muscle tropomyosin isoforms, including Tpm2.1, Tpm1.7, and Tpm3.1, reduce myosin 1 binding to actin^47^.

We first asked whether Tpm2.1 may negatively regulate GPI-AP nanoclustering (Fig. 3A). To test this, we used MDA-MB-231 cells, a human breast cancer cell line where the TPM2 gene is not expressed^48,49^. These cells are highly migratory, invasive and have been shown to be less sensitive to substrate stiffness^49,51^. The absence of TPM2 expression in these cells was confirmed by western blot and transcriptomic analysis (Supplementary Fig. S2A,B). We predicted that if Tpm2.1 limits actin accessibility to myosin 1, then the absence of Tpm2 would be expected to promote clustering, whereas re-expression of Tpm2.1 should suppress it. Consistent with this prediction, re-expression of WT Tpm2.1 in MDA-MB-231 cells reduced GPI-AP nanoclustering as confirmed by the increase in emission anisotropy of mRuby-GPI in MDA-MB-231 cells transfected with YFP-Tpm2.1 (Fig. 3B).

**Fig. 3:**
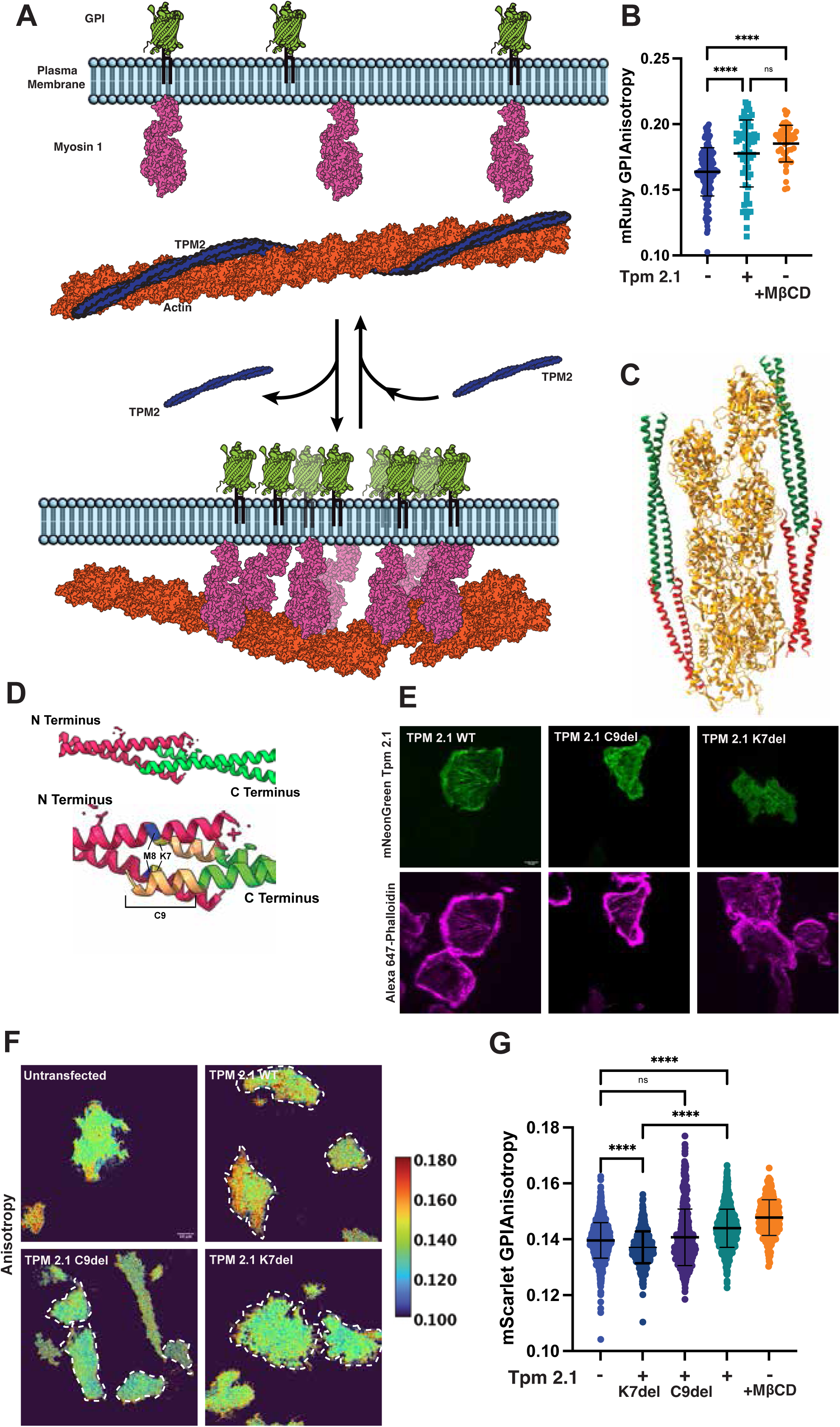
Tpm2.1 negatively regulates GPI-AP nanoclustering. (A) Schematic depicts gating of myosin1-actin interaction by tropomyosin wherein removal of tropomyosin permits myosin 1 interaction with actin, promoting GPI-AP-nanoclustering. B) Scatter plots show mean ±s.d of emission anisotropy determined from ROIs (2.2 x 2.2 um^2^) from cells expressing mRuby-GPI in MDA-MB-231 cells (blue), with YFP Tpm 2.1, (teal) and after MβCD treatment (orange), as indicated. Data shown from representative experiment, N=3. Each condition from >15 cells C) Redrawn EM structure of cardiac muscle thin filaments depicting Tpm head-tail interactions of two Tpm pairs (red and green) on actin. Structure adapted from PDB: 8DDO. TroponinT2 not shown for clarity. D) Schematic depicts Tpm2.1 N-C terminal dimers (top), highlighting the region of overlap between overlapping dimers (below) where some of the muscle myopathy mutants lie, and have been selected to disrupt Tpm-actin interactions. E) TIRF images show actin localisation (phalloidin Alexa 647, magenta) of mNeonGreen-tpm 2.1 and the indicated mutants (green). F) Anisotropy maps of mScarlet-GPI expressed in MDA-MB-231 cells alone or transfected with the indicated constructs. Pink dotted lines indicated cells transfected with the indicated Tpm constructs. G) Scatter plots show mean ±s.d of emission anisotropy determined from ROIs (2.2 x 2.2um^2^) of mScarlet-GPI expressed in MDA-MB-231 cells alone or co-expressed with mNeonGreen-tagged Tpm 2.1 (teal), K7del (dark blue) or C9del (purple), or treated with MβCD treatment (orange). Data shown from representative experiment, N=2. Each condition from > 26 cells. All intensity and anisotropy images were collected using TIRF emission anisotropy microscopy. *p* values determined from Kruskal-Wallis test with Dunn’s multiple comparisons test, where ns indicates no significant difference and **** indicates p <0.0001. Scale bar, 10um.

We next sought to perturb Tpm2.1-actin association more directly. Tropomyosin association with actin depends not only on direct tropomyosin-actin interactions but also on cooperative head-to-tail overlap between adjacent tropomyosin dimers^59^ (Fig. 3C,D). We reasoned that the disruption of the end-end association of two pairs of dimers could reduce the stability of Tpm 2.1 on actin. For these disruptions, we looked towards mutations commonly found in myopathies and mutations, some of which have also been shown to reduce actin association of skeletal and cardiac Tpm to actin filaments in *in vitro* reconstitution experiments^60^. We therefore generated two Tpm2.1 mutants predicted to destabilise actin filament association: an N-terminal lysine 7 deletion (K7del), based on a conserved residue implicated in nemaline myopathy, and a C-terminal deletion of the final nine amino acids (C9del), designed to disrupt the head-to-tail overlap region (Fig. 3D). Recombinant K7del skeletal Tpm/Tpm2.2 mutants are unable to incorporate normally into sarcomeres, and patients with this mutation show increased myofibre calcium sensitivity^60,61^. Mutations and deletion in the last exon of Tpm are also associated with mild to severe nemaline myopathy and cardiomyopathy, depending on the nature of the mutation^62–66^. Based on this we reasoned that a similar mutation in non-muscle Tpm 2.1 may also disrupt actin association.

Consistent with our reasoning, mNeonGreen-tagged K7del and C9del exhibited altered localisation relative to WT Tpm2.1 when expressed in MDA-MB-231 cells (Fig. 3E). Compared to wild type Tpm2.1 which exhibits a prominent localization on actin filaments, C9del and K7del both show inconsistent localization to actin filaments and exhibit a significant cytoplasmic distribution. We then examined GPI-AP nanoclustering in MDA-MB-231 cells expressing these constructs. While expression of WT Tpm2.1 produced a reduction in clustering evidenced by the increase in emission anisotropy of mScarlet-GPI, expression of either K7del or C9del mutants failed to elicit the same response. Indeed, clustering in these mutant backgrounds appeared to increase slightly relative to parental MDA-MB-231 cells (Fig. 3F,G). These findings are consistent with the idea that stable Tpm2.1 association with actin negatively regulates nanoclustering.

### Vinculin promotes nanoclustering by modulating tropomyosin-actin interactions

Since vinculin’s actin-binding activity is required for restoration of clustering, and as tropomyosin polymerises cooperatively along actin filaments, we hypothesised that vinculin binding to actin may influence tropomyosin occupancy on the filament. There is precedent for this type of regulation: actin-binding proteins such as fimbrin can displace tropomyosin from actin filaments, rendering the filament accessible to ADF/cofilin-mediated severing^67^. We therefore asked whether vinculin-tail binding to actin might similarly compete with or displace tropomyosin *in silico*.

Tropomyosin is known to form contacts with residues D25,K326, K328 and P333 closely juxtaposed to the actin filament^35,68–70^; the actin binding domain of vinculin also binds to residues 326,329,332 and 335^71–73^. These common binding sites suggest competition for vinculin and tropomyosin to bind actin filaments. To further examine this possibility, we compared the actin-binding surfaces of vinculin tail and tropomyosin using available structural assemblies of vinculin tail-actin (3JBI)^73^ and tropomyosin-actin (3J8A)^70^ (Fig. 4A-B). Because structural assemblies containing Tpm2.1 specifically are not currently available, we used actin-bound tropomyosin structures from related isoforms, including skeletal muscle tropomyosin 3J8A, (Fig. 4 B) 6KN7, 5NOG^74,75^ (Supplementary Fig. S3A, B), non-muscle Tpm1.6 7ZTC, and non muscleTpm3.2 7ZTD (Supplementary Fig. S3C, D). The analysis shown here is based on 3J8A. Structural superposition on actin filament suggests that vinculin tail and tropomyosin engage overlapping or closely adjacent surfaces on actin, consistent with the possibility of steric competition or displacement (Fig. 4C,D).

**Fig 4:**
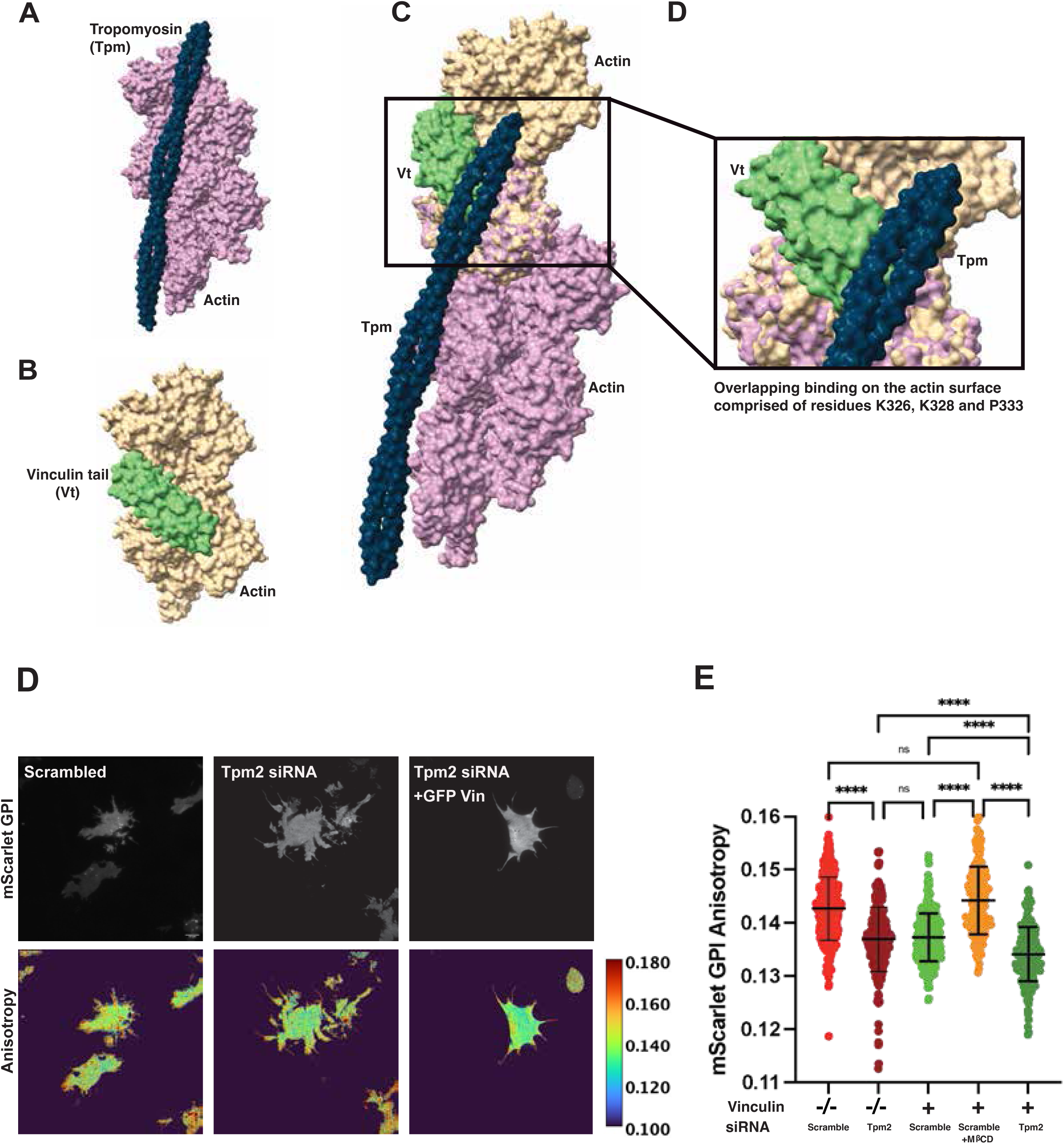
Vinculin promotes nanoclustering by modulating tropomyosin-actin interactions. A, B) Tropomyosin dimer (blue) on actin filaments (pink) redrawn from a x-ray structure (3J8A) in the PDB and corresponding vinculin tail (green)-actin filament (yellow) interaction structure (3JBI; B). C-D) Structural superposition of tropomyosin (from 3J8A) and Vinculin tail (from 3JBI) on the same aligned actin filament structure (RMSD between 356 pruned atom pairs = 1.042 Å, Across all 367 pairs = 1.128 Å) shows that they engage on overlapping surfaces. D) Intensity images and anisotropy maps of mScarlet-GPI expressed in MDA-MB-231 cells alone or transfected with the Tpm2 siRNA and GFP-Vin constructs. E) Scatter plots show mean ±s.d of emission anisotropy determined from ROIs (2.2 x 2.2um^2^) of mScarlet-GPI expressed in Vin -/- cells with 50nM scrambled control siRNA (red), or 50nM TPM2 siRNA (brown), or with GFP-Vin (green, dark green) with scrambled siRNA or TPM2 siRNA without or with MβCD treatment (orange). Structural superposition performed in UCSF ChimeraX. Data shown from representative experiment, N=2. Each condition from >25 cells. All intensity and anisotropy images were collected using TIRF emission anisotropy microscopy. *p* values determined from Kruskal-Wallis test with Dunn’s multiple comparisons test, where ns indicates no significant difference and **** indicates p <0.0001. Scale bar, 10um.

Based on this model, we predicted that if vinculin promotes myosin 1-dependent clustering by reducing tropomyosin occupancy on actin, then removal of tropomyosin in Vin-/- cells should restore clustering by increasing actin availability for myosin 1 even in the absence of vinculin. To test this, we depleted Tpm2 using siRNA in Vin-/- cells and examined GPI-AP nanoclustering. Tpm2 siRNA treatment depletes Tpm2 and other isoforms, whereas the scrambled siRNA sequence treatment does not alter the levels of Tpm in Vin-/- cells (Supplementary Fig. S2C, D). Consistent with this prediction, siRNA mediated Tpm knockdown restored GPI-AP nanoclustering in Vin-/- cells, whereas scrambled siRNA treatment had no effect. As expected, re-expression of vinculin in Vin-/-cells also restores GPI-AP clustering (Fig. 4D,E). These observations support the role of Tpm2 as a negative regulator of GPI-AP clustering and its modulation via the presence of activated vinculin.

## Discussion

The results presented here provide evidence for a model in which activated vinculin promotes plasma membrane GPI-AP nanoclustering by relieving tropomyosin-mediated restriction of myosin1-actin interaction (Fig. 5). We envisage the cascade of events depicted in the model in Fig. 5, downstream of engagement of Integrins by extracellular matrix ligands (*e.g.* fibronectin) leading to RhoA activation followed by the generation of formin-mediated actin filament nucleation^29^. While many of the elements of the model have been addressed previously, here we identify tropomyosin as a crucial negative regulator of this pathway gating the construction of GPI-AP-nanoclusters and resultant meso-scale domains via deploying activated Vinculin.

**Fig 5:**
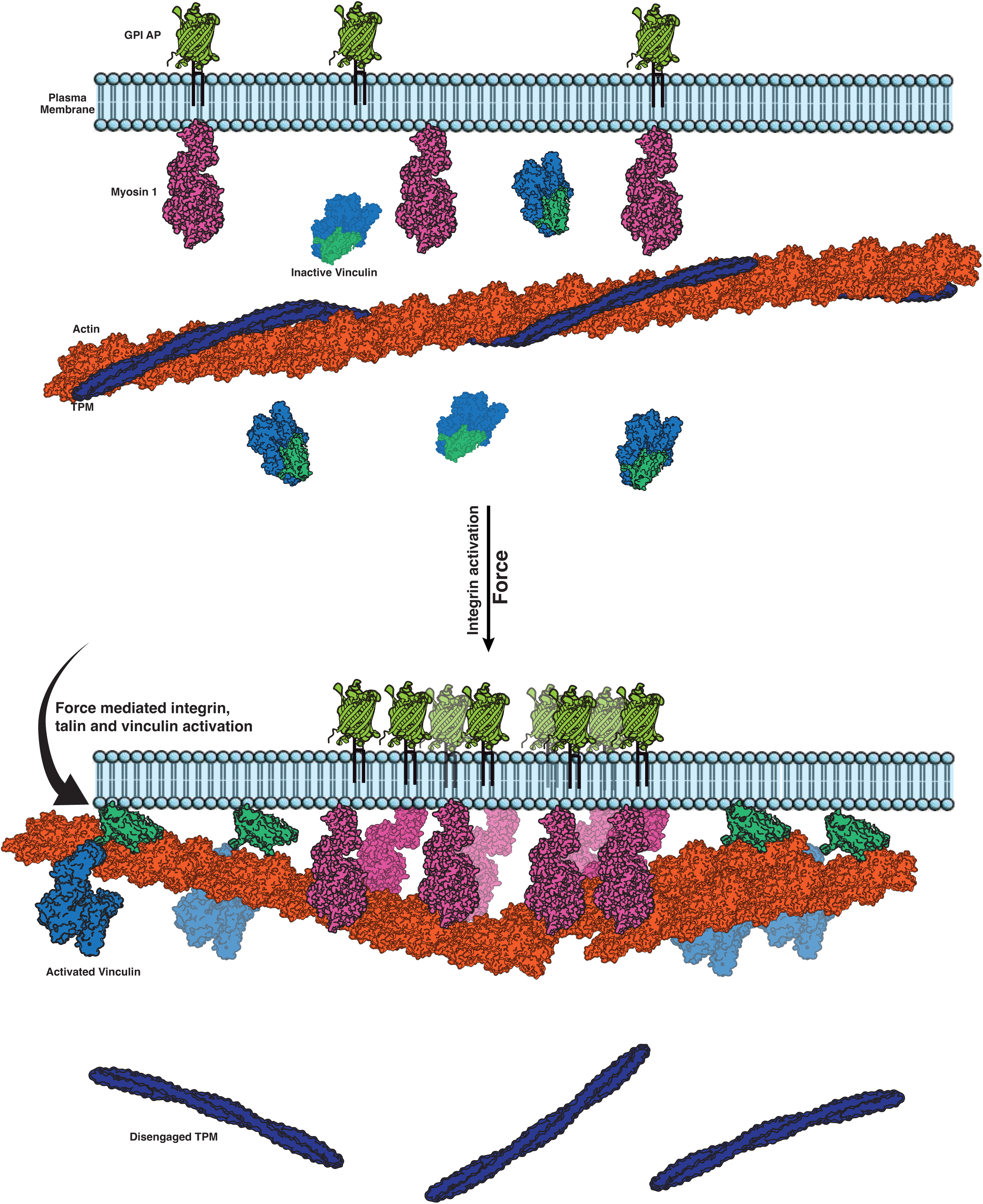
Model depicting vinculin-tropomyosin-myosin 1 mediated regulation of GPI-AP nanoclustering. Schematic depicts the proposed model for regulation of GPI-AP nanoclustering. In a quiescent state (for example in the absence of integrin engagement), vinculin remains in its auto-inhibited closed state with Tpm coating juxta-membrane actin filaments, limiting access to membrane associated myosin 1. Upon integrin activation, talin engagement leads to the opening of the vinculin molecule. This leads to the exposure of the vinculin tail with its actin and lipid binding sites, displacing tropomyosin from membrane proximal actin, and making actin available for myosin1-mediated GPI-AP nanoclustering.

Tropomyosins are key regulators of actin filament identity. They not only stabilize actin but also specify which motors and actin-binding proteins engage a given filament, sorting the actin cytoskeleton into functionally and spatially distinct populations^34^. A classic example of this is in muscle, where tropomyosin, together with troponin, gates access of myosin II to actin in a calcium-regulated manner. Although non-muscle actin networks lack the canonical troponin switch, the logic is similar: tropomyosin isoforms mark filaments with distinct functional identities by selectively permitting or excluding classes of motors and actin-binding proteins. It has been reported that while Tpm1.7 and 3.1 enhance non-muscle myosin 2’s ATPase activity, Tpm2.1 does not have this effect^47^. Non-muscle tropomyosins also regulate actin access and reduce ATPase activity of myosin 1C^47^. Our results suggest that this classical principle of tropomyosin-dependent motor gating observed at the sarcomere in muscle also applies at the plasma membrane of non-muscle cells, where Tpm2.1 appears to restrict access of myosin 1 to the actin filaments that promote GPI-AP nanoclustering.

This interpretation is supported by the behaviour of MDA-MB-231 cells. These cells lack TPM2 expression and have abundant GPI-AP nanoclusters; reintroduction of WT Tpm2.1 suppresses GPI-AP nanoclustering. By contrast, mutants (Tpm2.1C9del and Tpm2.1K7del) predicted to weaken stable tropomyosin filament assembly show markedly reduced localization to actin and fail to inhibit GPI-AP clustering. This indicates that the inhibitory effect depends on stable tropomyosin occupancy along actin filaments.

Our *in silico* structural superposition analysis suggest that the vinculin tail domain and tropomyosin engage overlapping or closely adjacent regions on actin filaments, raising the possibility that their association with actin may be mutually antagonistic. This provides a direct route by which a mechanosensitive adhesion protein such as vinculin could regulate actin filament availability. In muscle, tropomyosin-troponin complex gates myosin access to actin; in a non-muscle cells plasma membrane juxtaposed activated vinculin may locally reverse tropomyosin mediated actin decoration. This would then allow myosin 1 to engage membrane proximal actin filaments. In our analysis, vinculin tail’s association with actin seems to clash with all the isoforms and structures we assayed (Supplementary Fig. S3).

Together, these findings suggest that vinculin and tropomyosin define a mechanochemical switch that determines whether membrane-proximal actin can engage membrane-associated myosin 1, required to generate ordered, nanocluster-rich membrane domains. This may also explain how active-emulsion-like membrane domains could be spatially restricted during spreading and migration. Integrin engagement with fibronectin produces localized force, activating talin and vinculin at nascent adhesions. If activated vinculin locally reduces tropomyosin occupancy, generating myosin 1-permissive actin filaments, then the machinery required for GPI-AP nanoclustering would arise specifically at sites of mechanical engagement with the extracellular matrix. Ordered membrane domains would therefore not form uniformly across the cell surface, but at spatially defined cortical regions where force, adhesion signaling, and cortical actin state converge. These domains could then serve as local signaling platforms for lipid-modified signaling molecules such as Rac1, helping coordinate protrusion and spreading, necessary for durotaxis.

MDA-MB-231 cells lack Tpm2 and are highly invasive, raising the possibility that altered tropomyosin expression changes not only cytoskeletal mechanics but also the organization of membrane signaling platforms. If the loss of Tpm2.1 biases actin toward a myosin 1-permissive state, then transformed cells may gain an altered capacity to form active-emulsion-like membrane domains that support substrate-independent related-signaling such as observed in their anoiksis-resistance^49,50,76^. More broadly, differential expression of tropomyosin isoforms across developmental, mechanical, or pathological states could reshape membrane organization by tuning which actin populations are available for membrane-cortex coupling. This raises the possibility that membrane organisation itself is subject to regulation through changes in actin filament identity.

Our study supports a view of the plasma membrane as a mechanochemically regulated active emulsion, whose local organization emerges from the coupling of adhesion, force, lipid interactions, and actomyosin dynamics. In this framework, GPI-AP nanoclusters are not passive assemblies produced solely by lipid affinity or thermodynamic partitioning, but dynamic structures whose formation depends on the continuous input of mechanical and cytoskeletal activity. Together, our findings support a hypothesis where vinculin translates the mechanochemical activation of integrin into active plasma membrane organization by modulating the access of myosin 1 to membrane-proximal actin. This permits myosin 1 to engage actin, stabilize inner leaflet lipid organization, and drive transbilayer coupling to GPI-APs. Through these linked processes, localized mechanical inputs generate nanocluster-rich liquid-ordered membrane domains and thereby sustain the plasma membrane as an active emulsion. Vinculin and tropomyosin thus emerge as antagonistic regulators of a membrane–actin interface which integrates integrin activation, force, actin filament identity, and lipid organization into a dynamic system.

## Methods

### Cell culture

Vin-/- and Vin+/+ mouse embryonic fibroblasts ^31,32^ were maintained in low glucose DMEM (Invitrogen), supplemented with 1.8g/l NaHCO_3_, 25mM HEPES and supplemented with 10% heat inactivated FBS and PS-Glutamax (Invitrogen). MDA-MB-231 cells were maintained in high glucose DMEM, (Gibco) supplemented with 1.8g/l NaHCO3, 25mM HEPES and supplemented with 10% heat inactivated FBS and PS-Glutamax (Invitrogen).

### Plasmids, buffers and reagents

To generate mScarlet-GPI, mScarlet-I (Addgene, #85044) fragment was constructed via PCR with overhangs for insertion into the linearised mYFP-GPI backbone which carries the folate receptor secretory signal and GPI-anchor signal sequences (originally provided by M. Edidin). mNeonGreenHO-Tpm2.1 was mutated via PCR to delete K7 and the last 9 amino acids of the C-terminus. Human mNeonGreenHO-Tpm2.1 and human mYFP-Tpm2.1 were used for addback experiments in MDA-MB-231. GFP-Vt (Addgene, #46265) and GFP-Vin (*Gallus gallus*) were used for Vin addback in Vin-/- cells. Sequences for clones were verified by whole plasmid sequencing with Plasmidsaurus. Imaging was performed in 1x M1 buffer (150mM NaCl, 20mM HEPES, 5mM KCl, 1mM CaCl_2_ and 1mM MgCl_2._ Cells were fixed in 4% paraformaldehyde diluted from 16% in PBS, pH 7.2.

### Perturbations and drug treatments

10mM Methyl-β-cyclodextrin (MβCD) was prepared in 1x M1-glucose +0.5% FBS. Cells were incubated with MBCD for 45 mins, with a change of buffer every 15mins. Pentachloropseudilin ^53,54^ was reconstituted from powder to 50mM in DMSO. Cells were pre-treated with 2.5-3uM PClP for 1.5h, and cell spreading was allowed to progress in the presence of the drug. DMSO was used as a control in all experiments with drug treatments.

### siRNA treatment

TPM2 SmartPool siRNA (Dharmacon) and scrambled control were transfected into cells using JetPrime transfection reagent (Polyplus) according to the manufacturer’s instructions. 50nM siRNA was transfected during cell plating, and cells were allowed to grow for 48h. Media was then replaced, and siRNA transfection mix was added again to adhered cells. After 48h of this second transfection, cells were imaged. This protocol was followed since siRNA KD in Vin-/- cells was very inefficient over the standard transfection protocol of 72h. siRNA mediated knockdown was confirmed by western blot.

### Western Blotting

Samples were analysed by SDS-PAGE and western blot using a 12-20% gradient gel (BioRad). 20ug protein was loaded per lane. Knockdown samples, controls and housekeeping genes were run on the same gel. Following wet transfer to nitrocellulose membrane, the blot was cut (dotted lines) for antibody hybridisation. Following primary and secondary antibody incubation, the cut blot was reassembled and probed with Pierce SuperSignal West Pico (ThermoFisher). Mouse Anti-Tropomyosin TM311antibody (Sigma) was used at a dilution of 1:500. Mouse Anti-Vinculin (clone V284, Sigma) was used at a dilution of 1:2000. Rabbit Anti-β-Tubulin (CST) was used at a dilution of 1:2000. HRP tagged anti-rabbit and anti-mouse secondaries (CST) were used at a dilution of 1:4000. Protein levels were quantified using integrated density measurements done on Fiji^77^.

### Coverslip preparation

No1.5 glass bottom dishes (CellVis) or 22x22 mm no 1.5 glass coverslips (Ted Pella) were used for imaging. Coverslips were prepared by 15 mins sonication in pure ethanol at 50°C. Following this, coverslips were allowed to dry in a laminar flow hood. Human plasma fibronectin (Merck) was diluted to 10ug/ml in PBS pH 7.2, and dishes or coverslips were incubated with the solution for 90 mins at 37°C or 4°C overnight. After coating, excess solution was removed, and dishes/coverslips were washed gently with PBS twice. After washing, dishes/coverslips were immediately used for cell spreading assays.

### Cell spreading assays

Cells were serum-starved in incomplete DMEM medium (without serum supplementation) for 3h prior to the assay. Starved cells were de-adhered using TryplE (Invitrogen) and resuspended in pre-warmed spreading medium (DMEM incomplete medium+0.5% FBS). Cells were centrifuged at 100xG, and the pellet was resuspended in 1.5ml spreading medium. This was added to fibronectin-coated coverslips. Cells were allowed to adhere at 37 for 10mins, after which any unadhered cells were washed off. Cells were then allowed to spread at 37°C for 1-1.5hrs, then taken for further imaging.

### Phalloidin labelling

Coverslips/dishes with cells spreading were fixed with 4% PFA in PBS for 20mins at RT. PFA was then removed and cells were permeabilized in 0.1% TritonX+PBS for 5mins. Following this, they were incubated with Alexa 647-Phalloidin (Invitrogen) for 1h. Excess phalloidin was removed by 3x PBS wash, and fixed and labelled cells were stored in 1xPBS for up to 1 week.

### Live cell microscopy

Live cells were imaged in M1 imaging buffer on an inverted Nikon TiE microscope with a 100x 1.49NA objective based TIRF mode with polarized excitation passing through a clean-up polarizer to ensure high extinction ratio. The system was equipped with two Prime95B sCMOS cameras (Photometrics), imaged simultaneously using a Cairns splitter. Emission anisotropy was measured by splitting emission into respective parallel and perpendicular polarized components using a nanogrid polarizing beam splitter (Moxtek FB04C) as described^52^. mScarlet and mCherry fluorophores were excited using a 561nm laser, and emission was collected through a quadband dichroic (Chroma ZT405/488/561/640rpcv2) and a 600/50 emission filter. mYFP and EGFP were imaged with a 488nm laser and emission was collected through a 488 DM and 525/50 emission filter. Alexa 647 labeled probes were imaged using a 647nm laser. Images were acquired using Micromanager V2.0.0. G factor images for anisotropy measurements were obtained using appropriate dyes (FITC <10uM in water for 488nm channel imaging and Rhodamine-6-G <10uM for 561nm channel imaging). Dark images for background correction were acquired at the same laser power and integration time using imaging buffer.

### Analysis

In brief, raw fluorescence microscopy images for PA and PE channel images were converted to photoelectrons. Pixel registration was performed to align PE channel images to PA channel images, and dark images were subtracted to remove image background. G-factor correction was applied to correct for any polarisation bias in the system. These corrected images were used to draw 2.2x2.um ROIs from which mean pixel intensity was extracted using ROI manager in Fiji image analysis software. Anisotropy was calculated using the following formula:

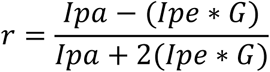

ROIs were drawn on flat, feature free membrane regions of the cells, and regions with low photons counts and intensity features were avoided to prevent noise or membrane feature related artefacts. Calculated anisotropy from ROIs in each condition was plotted as scatter plots, and statistical analyses were performed using GraphPad Prism 10 software. Sample sizes and statistical tests/significance are mentioned in respective figure legends.

### Structural overlap analysis

Structural overlap analysis was done using the Matchmaker tool in UCSF ChimeraX 1.9. PDB structures were imported in Chimera and analyzed using the Matchmaker tool. Resulting superimpositions were exported as image files. Alignment scores and RMSD are mentioned in respective figure legends.

### RNA sequencing

RNA sequencing was performed by Plasmidsaurus using Illumina Sequencing Technology with custom analysis and annotation.

## Acknowledgements

We thank Wolfgang H. Goldmann (Friedrich-Alexander-Universität Erlangen-Nürnberg, Erlangen,Germany) and Klaus M. Hahn (UNC-Chapel Hill School of Medicine Chapel Hill, NC, USA) for Vin-/- and Vin +/+ cells. We thank Peter W Gunning (UNSW, Sydney Australia), Will Scott and Mohan Balasubramanian (Warwick Medical School, University of Warwick, Coventry,UK) for YFP Tpm2.1 and mNeonGreen TPM constructs. We thank Parvinder Pal Singh (Indian Institute of Integrative Medicine (CSIR), Canal Road, Jammu, India) for providing PClP. We thank Gayathri Pananghat (IISER, Pune,India) for helpful discussions regarding structural analysis. We thank all members of the Mayor laboratory, especially Shubhangi Sharma for discussions regarding the study and for their comments on the manuscript. We thank the NCBS Central Imaging and Flow Facility and Greeshma Pradeep S and Matthew Gleeson from Mayor lab for help with optical setups. S.B acknowledges doctoral fellowship support from National Centre for Biological Sciences, Tata Institute for Fundamental Research. S.M. acknowledges support from Department of Biotechnology – Wellcome Trust India Alliance Margadarshi Fellowship IA/M/15/1/502018 and the Department of Atomic Energy, India (under Project No.RTI 4006), JC Bose National Fellowship (Department of Science and Technology, India), and S.M. and T. M.-W., Leverhulme Trust, UK (LIP-2021-017).

## Author contributions

S.B. and S.M. designed the study. Plasmid cloning and SDMs were done by S.B and T.M.-W. Cell culture, imaging and analysis were performed by SB. SB and S.M wrote the paper.

## Notes

### Competing Interest Statement

The authors have declared no competing interest.

