## Supplementary Material for "Tropomyosin and Vinculin antagonistically mediate GPI-anchored protein nanoclustering"

**Supplementary materials contain:**

Supplementary Fig. S1
Supplementary Fig. S2
Supplementary Fig. S3

Fig. S1

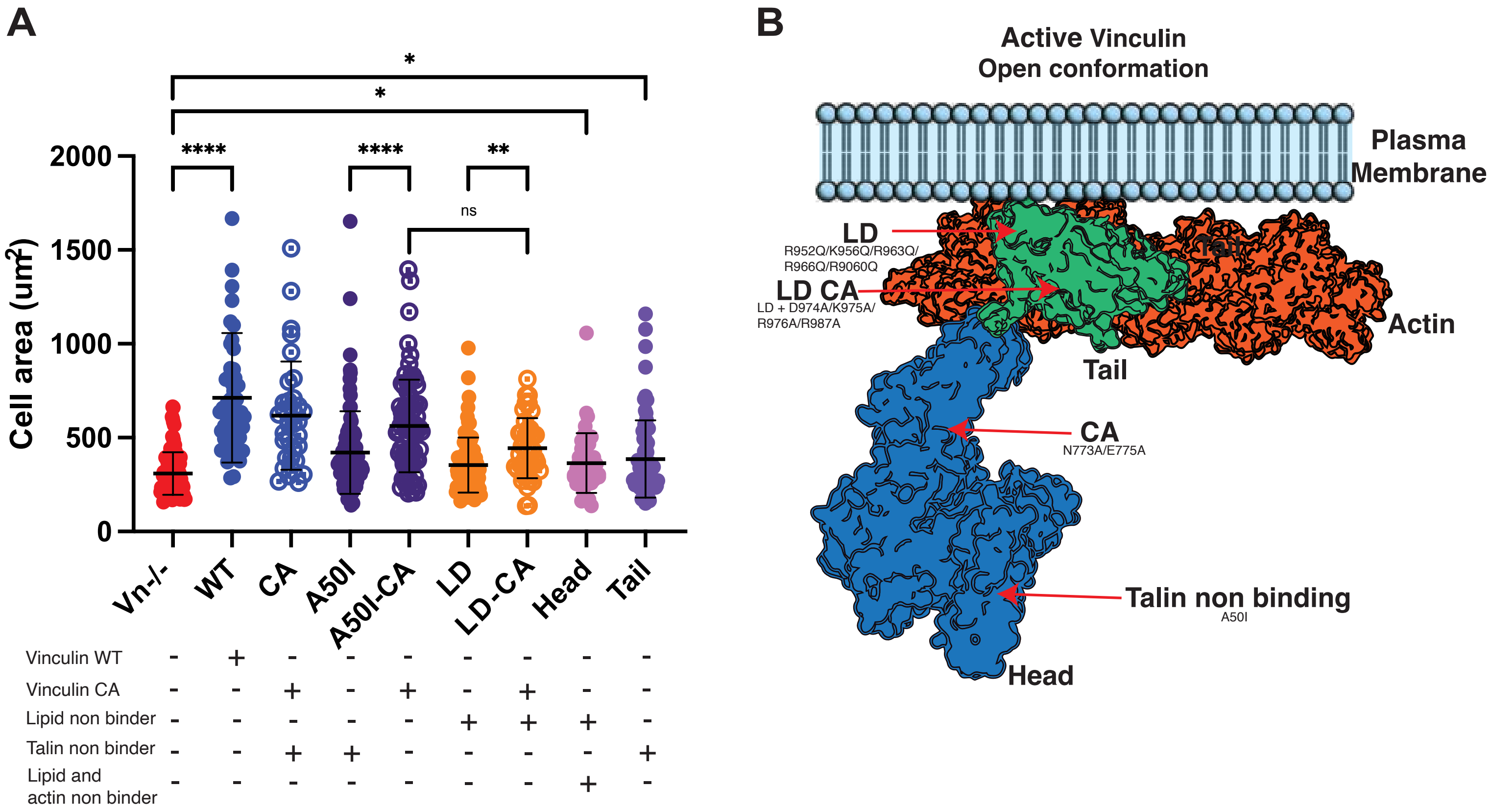

A
*
*
****
****
**
ns
Cell area (um²)
2000
1500
1000
500
0
Vn-/-
WT
CA
A50I
A50I-CA
LD
LD-CA
Head
Tail
Vinculin WT - + - - - - - - -
Vinculin CA - - + - + - + - -
Lipid non binder - - - - - + + + -
Talin non binder - - + + - - - - +
Lipid and actin non binder - - - - - - - + -
B
Active Vinculin
Open conformation
Plasma Membrane
LD
R952Q/K956Q/R963Q/
R966Q/R9060Q
LD CA
LD + D974A/K975A/
R976A/R987A
Tail
Actin
Tail
CA
N773A/E775A
Talin non binding
A50I
Head

**Supplementary Fig. S1**: **Vinculin mutants influence cell spreading** A) Scatter dot plots of area measurements of vinculin mutant addbacks in Vin-/- cells. Key underneath the plot depicts functions of each mutant. B) Schematic depicts sites for various vinculin mutants. N=2. Each condition from >16 cells *p* values determined from Kruskal-Wallis test with Dunn's multiple comparisons test, where ns indicates no significant difference and **** indicates *p* <0.0001

Fig. S2

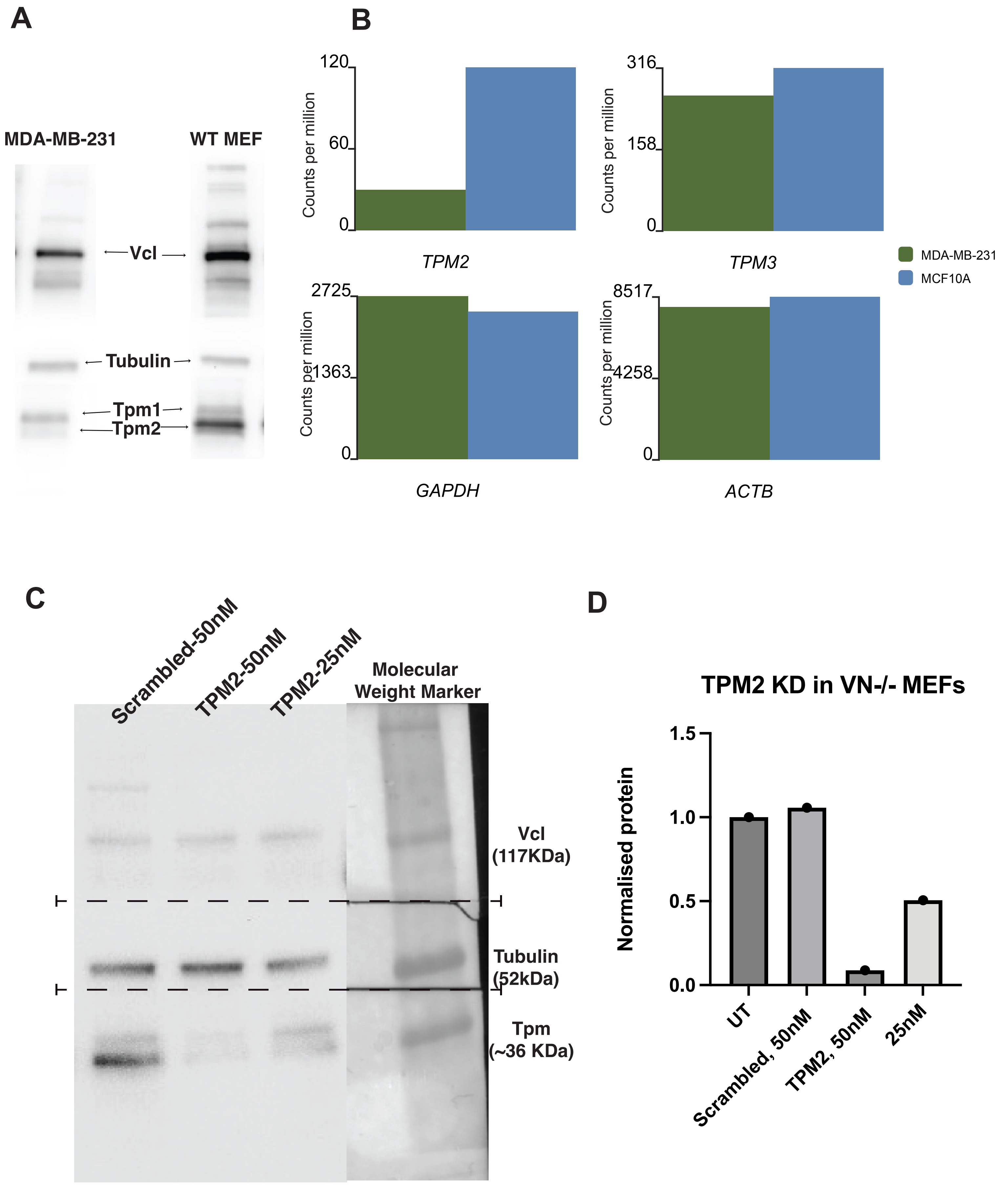
A
B
MDA-MB-231
WT MEF
Vcl
Tubulin
Tpm1
Tpm2
120
60
0
Counts per million
TPM2
316
158
0
TPM3
2725
1363
0
GAPDH
8517
4258
0
ACTB
MDA-MB-231
MCF10A
C
Scrambled-50nM
TPM2-50nM
TPM2-25nM
Molecular Weight Marker
Vcl (117KDa)
Tubulin (52kDa)
Tpm (~36 KDa)
D
TPM2 KD in VN-/- MEFs
Normalised protein
1.5
1.0
0.5
0.0
UT
Scrambled, 50nM
TPM2, 50nM
25nM

### Supplementary Fig. S2: Tpm2 expression levels

A) Western blot showing loss of high molecular weight Tpm2 from MDA-MB-231 cells. Tpm2 band indicated in MEFs for comparison. B) RNA sequencing data from MDA-MB-231 cells and from MCF10A cells. Bar plots depict counts per million for transcripts of Actin (ACTB), GAPDH(GAPDH), Tpm3 (TPM3) and Tpm2 (TPM2). Actin, GAPDH and Tpm3 represented as controls. C) Western blot showing the reduction of Tpm2 upon siRNA treatment with the loading control for the scrambled siRNA and different concentrations of Tpm2 siRNA. D) Bar graph shows levels of Tpm2 expression in Vin-/- MEF cells that were treated with siRNA against Tpm2. Protein levels for Tpm2 were normalized to the scrambled control.

Fig. S3

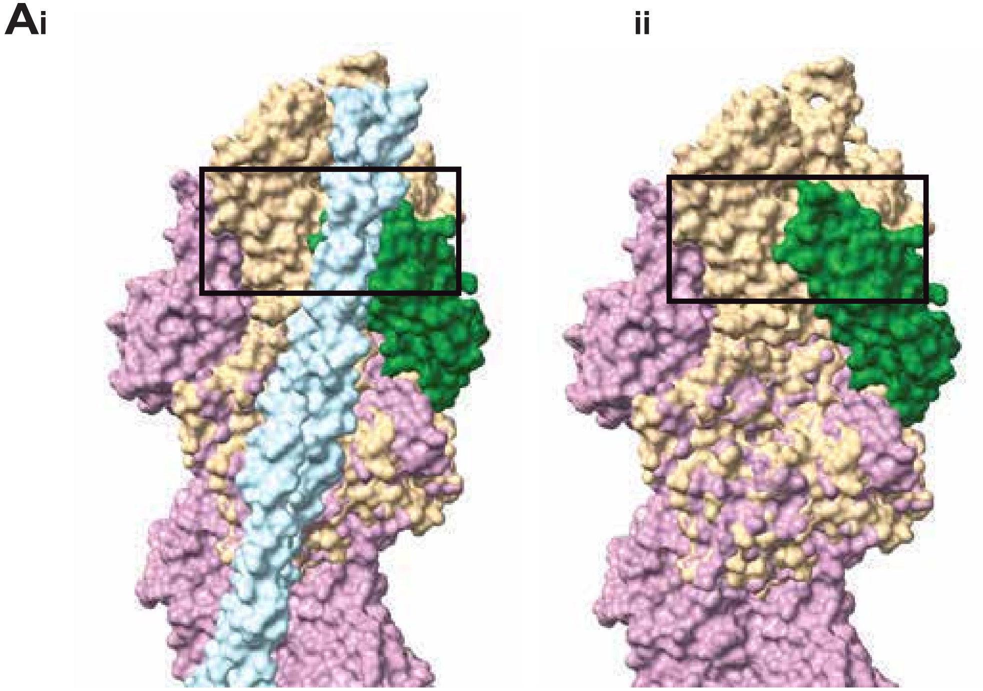


6KN7:Human cardiac thin filament in the calcium free state & 3JBI: Vinculin tail- Actin complex

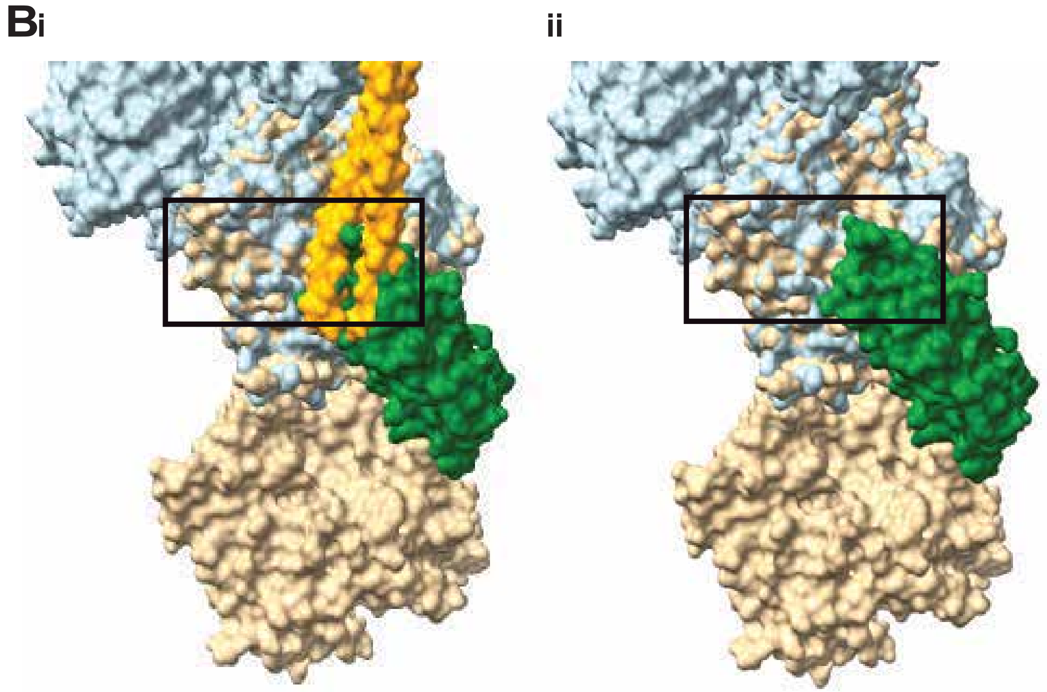


5NOG: Native Cardiac Thin Filaments - "Blocked" state & 3JBI: Vinculin tail - Actin complex

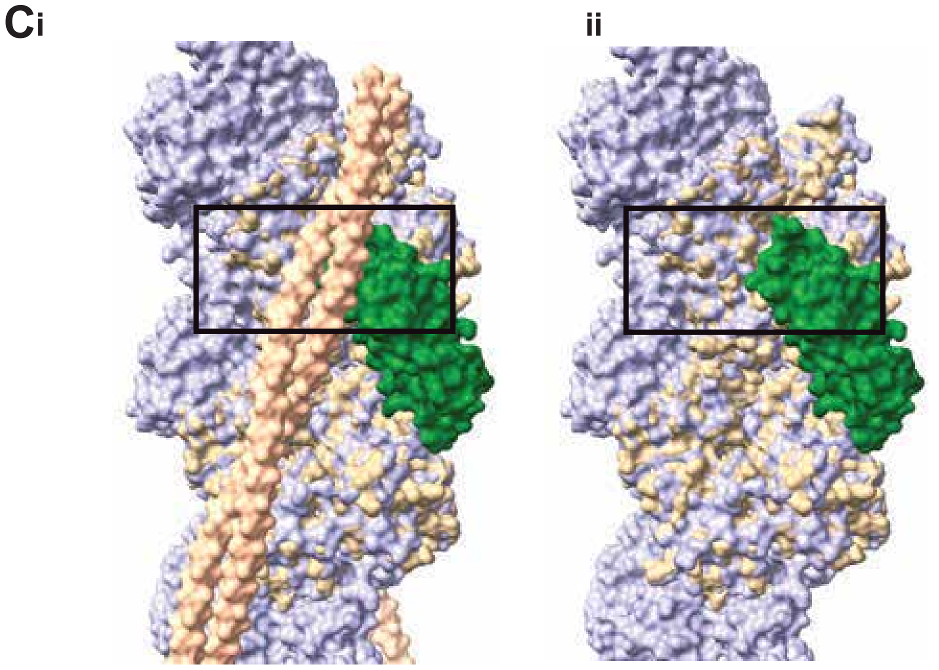


7ZTD:Non-muscle F-actin decorated with non-muscle tropomyosin 3.2 & 3JBI: Vinculin tail- Actin complex

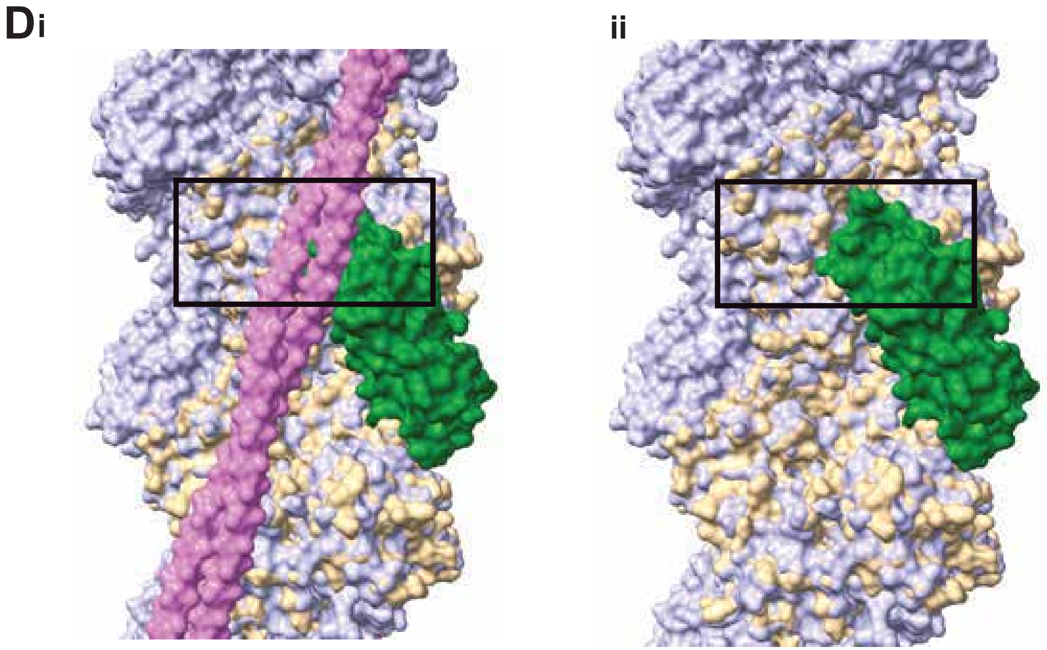


7ZTC:Non-muscle F-actin decorated with non-muscle tropomyosin 1.6 & 3JBI: Vinculin tail- Actin complex

**Supplementary Fig. S3: Superposition of tropomyosin isoforms and vinculin on structurally aligned actin filaments.** Vinculin tail domain (green) on actin filament (tan) from PDB structure, 3JBI was superimposed on cardiac muscle tropomyosin filament (blue) draped over an actin filament (pink) from PDB structure 6KN7 (A), or on tropomyosin (orange) and actin (blue) from cardiac muscle PDB structure 5NOG (B), or non-muscle Tpm3.2 (peach) and actin (lavender) structure from 7ZTD (C) or non-muscle Tpm1.6 (pink) and actin (lavender) from 7ZTC (D). Structures of troponin where available are deliberately hidden from view for the purpose of clarity. Matchmaker sequence alignment score for A = 1727.3. RMSD between 296 pruned atom pairs is 1.195 angstroms; across all 367 pairs: 1.739; B = 1631.9. RMSD between 331 pruned atom pairs is 1.156 angstroms; across all 367 pairs: 1.397; C = 1725.1 RMSD between 349 pruned atom pairs is 1.014 angstroms; across all 365 pairs: 1.149; D= 1700.5 RMSD between 344 pruned atom pairs is 0.937 angstroms; across all 365 pairs: 1.131. In all sub-sections ii) superposition with tropomyosin is hidden to reveal the binding footprint overlap of the Vinculin tail domains on the actin filaments. Black box denotes region of overlap. Note the vinculin tail and all Tpm isoforms are expected to bind to overlapping regions on the actin filament.
